# Comparative Unfolding of the Trp-cage Miniprotein in Anionic and Cationic Surfactants

**DOI:** 10.64898/2026.04.08.717321

**Authors:** Osita Sunday Nnyigide, Haewon Byeon, Uchenna Esther Okpete

**Author notes:** **Corresponding author:** Haewon Byeon.

## Abstract

The conformational dynamics of a model cationic protein in water and in the presence of anionic sodium dodecyl sulphate (SDS) and cationic cetyltrimethylamonium bromide (CTAB) surfactants at different concentrations were investigated using all-atom molecular dynamics simulations. Free-energy landscapes constructed along principal components reveal a compact, well-defined native basin at 25 °C in water, whereas elevated temperature (100 °C) induces a broadening of the conformational space and the emergence of multiple metastable states. The presence of surfactants further modulates this behavior in a concentration-dependent manner. Cluster population analysis shows that SDS promotes a highly heterogeneous ensemble characterized by reduced dominance of the native-like cluster, while CTAB partially protects the protein from thermal denaturation at higher concentrations. Radial distribution functions demonstrate strong accumulation of SDS headgroups around the protein and pronounced insertion of SDS alkyl tails into hydrophobic protein regions, indicating direct hydrophobic destabilization and micelle-assisted unfolding. In contrast, CTAB exhibits weaker headgroup association owing to electrostatic repulsion and reduced tail–hydrophobic contacts, suggesting a less disruptive interaction mechanism. At high concentration, CTAB aggregates provide a structured hydrophobic environment that stabilizes the folded state and suppresses denaturation. Together, these results provide a molecular-level picture of how surfactant chemistry and concentration govern the conformational stability of a cationic protein, highlighting the dominant role of hydrophobic interactions in surfactant-induced denaturation at high temperature.

**Graphical Abstract:** 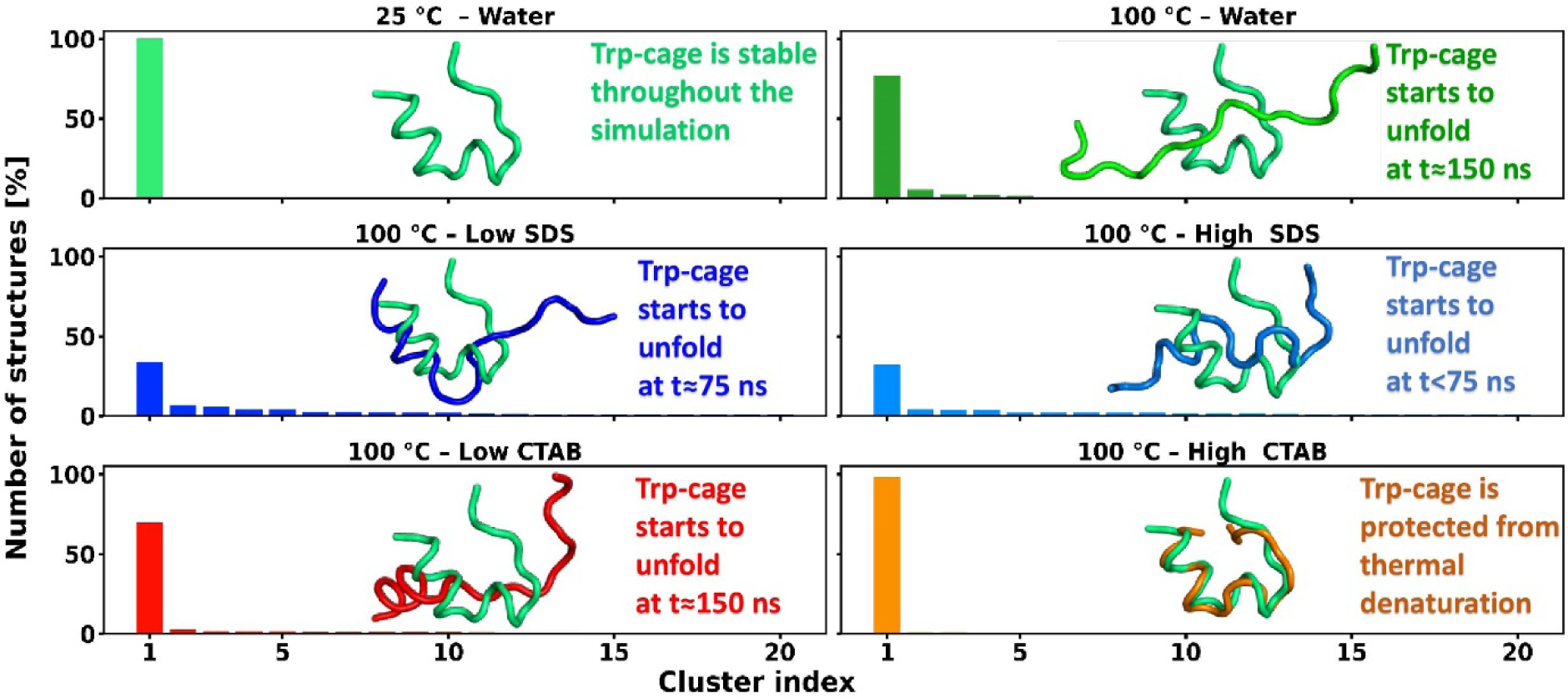

## 1. Introduction

Proteins are dynamic biomolecules whose biological activity depends critically on the preservation of their native three-dimensional structures [1–8]. Even subtle alterations in folding can drastically influence enzymatic activity, stability, solubility, and intermolecular interactions [2,3,7]. Therefore, understanding the factors that promote or inhibit protein unfolding remains a central theme in biochemistry, biophysics, and molecular biology. Among the numerous agents known to destabilize proteins—such as temperature, pH, denaturants, salts, and mechanical stress—surfactants represent a particularly important class because of their dual roles in both biological systems and industrial applications [9–16]. Surfactants are integral components of pharmaceutical formulations, detergents, food products, and nanomaterial systems, yet their interactions with proteins are complex, often non-linear, and highly dependent on surfactant charge and structure [17–22].

Protein–surfactant interactions are primarily governed by electrostatic attraction or repulsion, hydrophobic association, and cooperative binding events that differ significantly between anionic, cationic, and nonionic surfactants [3,7,12,17]. When surfactants bind to proteins, they can perturb secondary and tertiary structures, promote partial unfolding or molten globule states, and in some cases lead to complete denaturation [13,15]. The extent of unfolding does not depend solely on surfactant concentration but also on micellization behavior, headgroup charge, tail length, and the intrinsic stability of the protein itself [2,7,12]. Despite decades of research, the mechanisms by which oppositely charged surfactants induce unfolding remain an area of active investigation, especially when considering how charge polarity influences binding stoichiometry, molecular interactions, and the thermodynamics of unfolding.

The Trp-cage miniprotein serves as an excellent model system for probing these phenomena. Trp-cage is an ultrafast-folding, globular peptide consisting of 20 amino acids that adopts a well-defined tertiary structure centered around a hydrophobic core formed by a single tryptophan residue [16, 23]. Its structure is stabilized by a network of hydrophobic interactions, hydrogen bonds, and salt bridges, resulting in a compact α-helical fold. Owing to its small size, rapid folding kinetics, and well-characterized energy landscape, Trp-cage has become one of the most extensively studied systems in protein folding and stability research, particularly in computational and biophysical studies [16, 23]. Numerous experimental and simulation-based investigations have examined its folding–unfolding behavior under varying temperatures, solvent conditions, and in the presence of chemical denaturants [16,24,25]. Although Trp-cage lacks disulfide bonds and has a simpler architecture than larger proteins, its charged residues and exposed hydrophobic surfaces enable meaningful interactions with surfactants, making it a useful minimalist model for studying protein–surfactant interactions.

Anionic surfactants such as sodium dodecyl sulfate (SDS) are widely recognized for their strong denaturing power [26,27]. SDS binds cooperatively to proteins, initially associating with positively charged regions and subsequently disrupting hydrophobic cores once a critical binding threshold is reached [3]. These interactions often lead to pronounced structural rearrangements, including loss of α-helix content, expansion of the hydrodynamic radius, and destabilization of tertiary contacts [15,19]. Conversely, cationic surfactants such as cetyltrimethylammonium bromide (CTAB) exhibit different interaction profiles due to charge repulsion with the positively charged Trp-cage surface. As a result, CTAB may require specific concentration regime to induce significant unfolding, may stabilize certain intermediate states, or may promote partially unfolded structures that differ fundamentally from those induced by SDS. The contrasting behavior of anionic and cationic surfactants underscores the importance of electrostatic interactions as well as the subtle contributions of hydrophobic forces.

Although numerous studies have investigated protein–surfactant interactions individually, a direct and systematic comparison of Trp-cage unfolding in oppositely charged surfactants remains relatively limited [6,8,9,15]. Many previous works focus on a single surfactant type, or do not account for differences in binding cooperativity. This knowledge gap makes it challenging to fully understand how charge polarity alters not only the extent of unfolding but also the unfolding pathway, stability of intermediate states, and the underlying molecular mechanism.

Therefore, in this study, we conduct a comprehensive comparative analysis of the unfolding behavior of the Trp-cage miniprotein in representative anionic and cationic surfactants. Using computational approaches, we evaluate the thermodynamic features of unfolding, identify distinct intermediate states, and characterize the mechanisms responsible for structural destabilization. The findings offer new insight into the roles of surfactant charge, concentration, and molecular architecture in protein unfolding, thereby contributing to a deeper and more mechanistic understanding of protein–surfactant interactions.

## 2. Molecular dynamics simulation protocol

### 2.1 Preparation of the Protein and surfactant structures and co-ordinate files

The Trp-cage miniprotein (PDB ID: 1L2Y) [23] was obtained from the protein data bank website (https://www.rcsb.org) [28] which represent well-defined structure of the protein. SDS and CTAB structure files (PubChem CIDs: 3423265 and 3423265, respectively) were obtained from the PubChem website (https://pubchem.ncbi.nlm.nih.gov). The protein structure was checked for consistency using the PRAS server (https://www.protein-science.com/) [29]. The protonation states of the charged amino acid residues at physiological pH were maintained. The GROMACS topology files for the proteins were generated using GROMACS software (Van Der Spoel et al. 2005) [30] but for the surfactants, the topology files were generated automatically using the CGenFF software [31].

### 2.2 Final molecular dynamics production

Molecular dynamics simulations were performed in pure water and surfactant solutions using a cubic box type. The number of surfactant and water molecules required for each concentration were calculated following previous studies [15, 19]. The final surfactant concentrations were 0.28 M and 0.55 M (SDS or CTAB), hereafter referred to as the low and high concentrations, respectively. All simulations were performed at 0.15 M salt concentration to mimic physiological condition. The GROMACS open source software was used [30] and the interactions were described using the CHARMM36 force field [32] and TIP3P water model. Periodic boundary conditions were applied in all three directions of the box. The distance between the solute and the box was 1 nm. The short-range non-bonded interactions were calculated by applying 1.2 nm cut-off radii, while Particle Mesh Ewald (PME) method was used for long-range electrostatics calculation. Energy minimization of the system was performed using the steepest descent method with a total of 500, 000 steps at 2 fs time step. The simulations were performed for the systems at 25 and 1000 °C in the NVT- and NPT-canonical ensembles using velocity rescaling temperature coupling thermostat and Parrinello-Rahman barostat pressure coupling to reproduce correct kinetic ensembles. All heavy atoms were position restrained with the force constant of 1,000 kJ/mol nm^2^ for proper penetration/distribution of solvent. The LINCS algorithm [33] was applied to constrain all bonds involving hydrogen atom. The GRID method was used to search and update neighbour list with a frequency set to 10 steps. A total of 200, 000 steps were used at 2 fs time step for both temperature and pressure equilibrations. We plotted the total potential energy and total pressure to ensure that our system reached desired equilibrium conditions. The final MD production runs were carried out for 300 ns using leap-frog integrator. A time step of 2 fs was used. The trajectory was stored every 2 ps.

## 3. Results and Discussion

### 3.1. System Stability and Simulation Quality

To ensure that the observed structural changes in the Trp-cage arise from physical interactions rather than numerical artifacts, we first assessed the stability and equilibration quality of all molecular dynamics simulations. Systems were simulated in pure water at 25 °C and 100 °C, as well as in the presence of anionic (SDS) and cationic (CTAB) surfactants at 100 °C. Although the primary focus of this study is protein unfolding at elevated temperature and in the presence of surfactants, the simulation performed in pure water at 25 °C provides a baseline reference for the stable native structure of the Trp-cage protein.

The time evolution of the potential energy for all systems (Fig 1) shows a stable fluctuations around well-defined mean values, indicating proper equilibration. No systematic energy drift was observed over the production runs, confirming the numerical stability of the simulations. Temperature profiles remained tightly centered around the target values of 25 °C and 100 °C, respectively, with only small thermal fluctuations (Fig 2). Pressure fluctuations were similarly well behaved (data not shown), consistent with correctly functioning thermostat and barostat coupling. It should be noted that the observed fluctuations are due primarily to the protein’s small size and lack of extensive tertiary constraints. Such behavior is expected for miniproteins and ultrafast-folding peptides and does not indicate structural instability or simulation artifacts [16].

**Fig 1.**
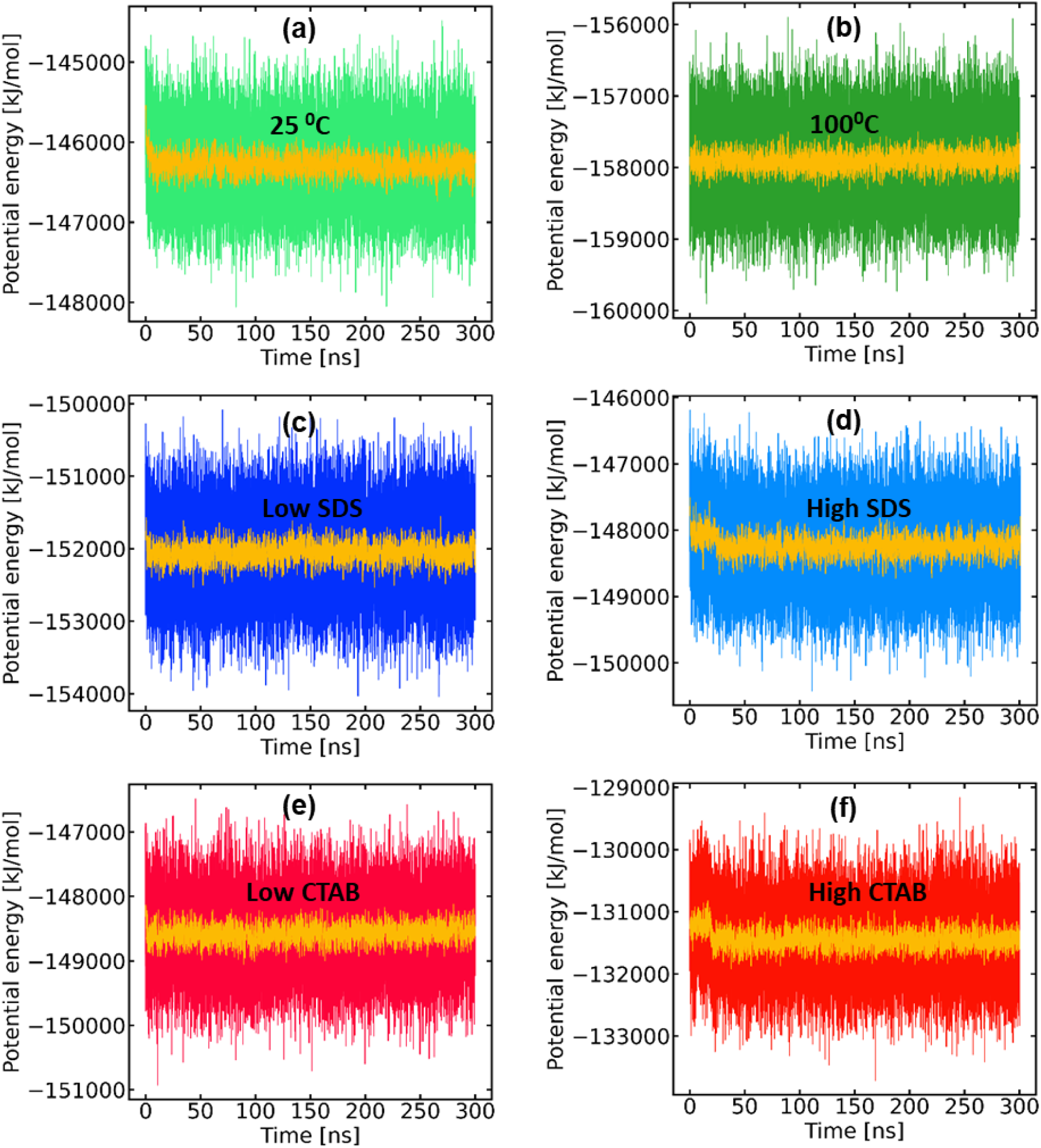
(a - f) Evolution of the potential energy of the systems and running averages.

**Fig 2.**
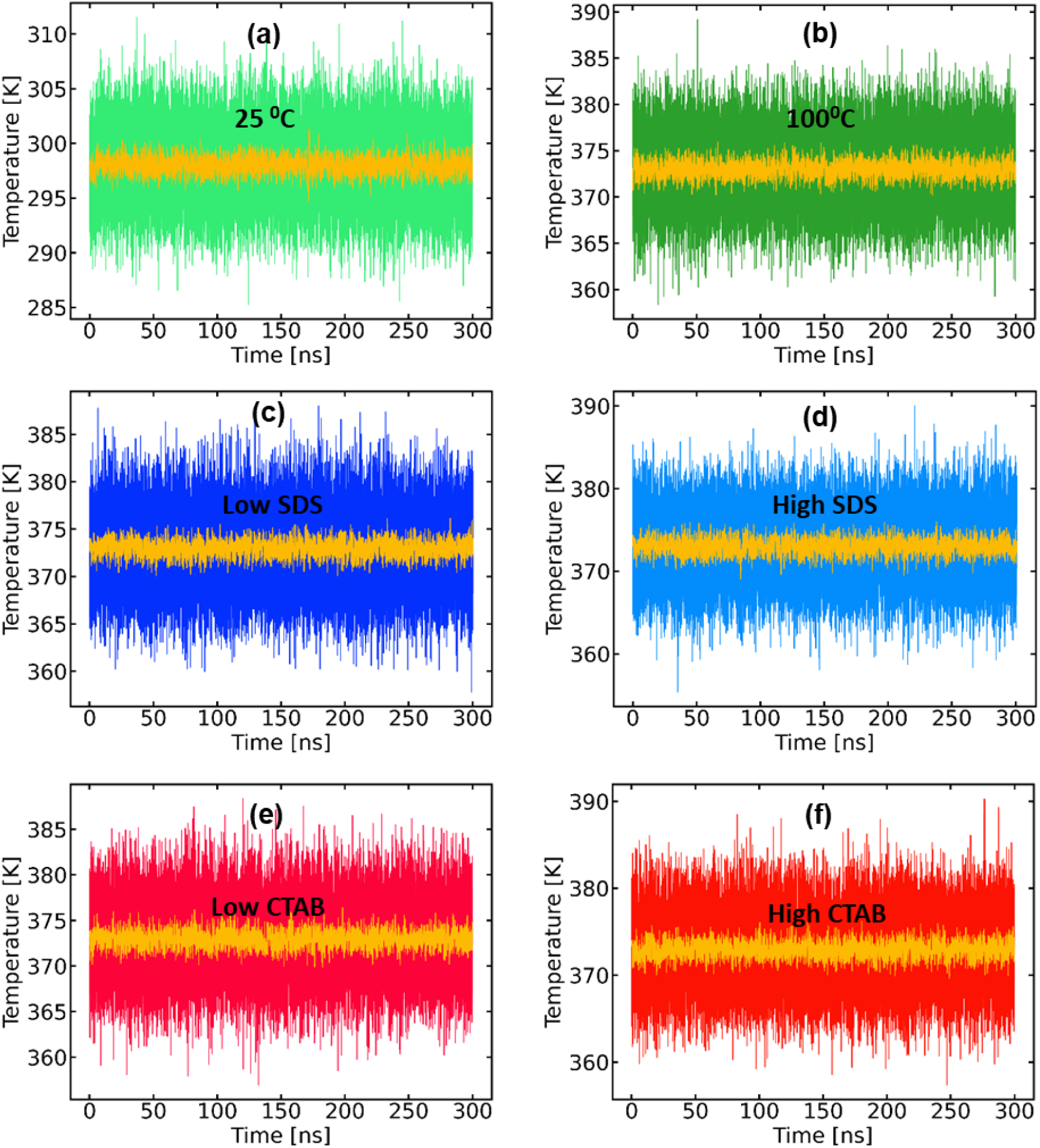
(a - f) Evolution of the temperature of the systems and running averages.

### 3.2. Root Mean Square Deviation Relative to the Native Structure

The structural stability of the Trp-cage miniprotein was evaluated by monitoring the behavior of the backbone root-mean-square deviation (RMSD) relative to the native structure in pure water and in the presence of surfactants [16, 17]. In pure water at 25 °C, the protein maintains a low RMSD throughout the simulation, with fluctuations characteristic of normal thermal motion in a small miniprotein [16] (Fig 3). At elevated temperature, unfolding in water is delayed and heterogeneous, indicating that temperature alone is insufficient to rapidly disrupt the native fold on the simulated timescale (i.e., the RMSD remains low up to 150 ns for pure water at 100 °C, indicating delayed unfolding).

**Fig 3.**
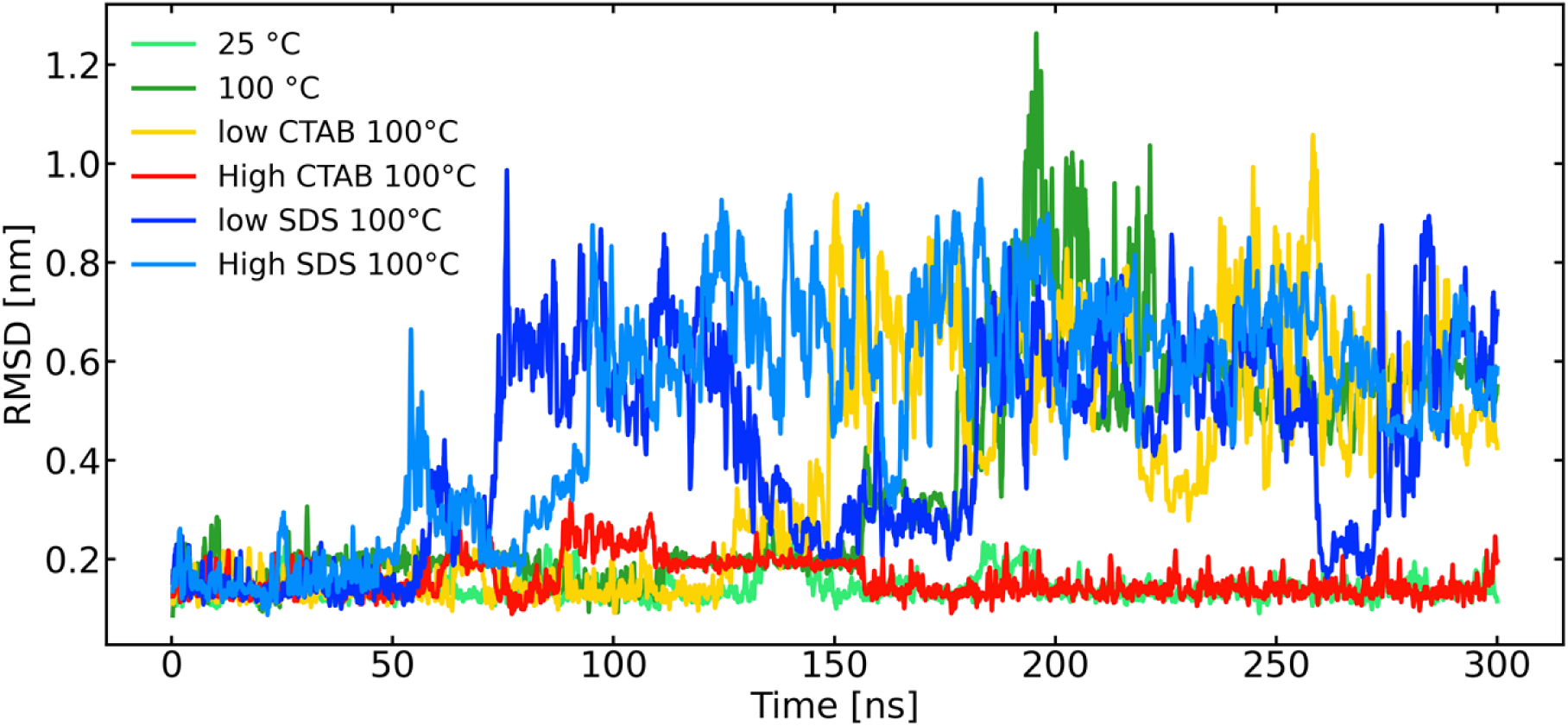
Evolution of the backbone RMSD of trp-cage on the simulated timescale (running average).

In contrast, simulations performed at 100 °C in the presence of surfactants display markedly different behavior. The SDS system shows an earlier and more pronounced increase in RMSD, with no clear plateau over the same timescale, consistent with progressive loss of native structure. The CTAB system exhibits a delayed and less extensive RMSD increase, indicating partial destabilization but greater structural retention compared to SDS. Comparatively, CTAB exhibits a markedly weaker destabilizing effect on Trp-cage. At lower CTAB concentration, the RMSD remains low up to 120 ns, suggesting partial unfolding. At higher CTAB concentration, the backbone RMSD remains low and comparable to that observed in pure water at 25 °C, indicating preservation of the native fold even under high-temperature conditions. This behavior can be attributed to electrostatic repulsion between the positively charged Trp-cage and the cationic CTAB headgroups, which limits cooperative binding and delays surfactant-induced destabilization. Consequently, CTAB primarily perturbs surface regions of the protein and stabilizes partially unfolded intermediates rather than inducing the extensive unfolding observed for anionic SDS. These results highlight the critical role of surfactant charge polarity in determining both the extent and pathway of Trp-cage unfolding [3, 7, 12, 15, 19].

While RMSD provides a measure of structural deviation from the native state, the radius of gyration [16] and DSSP [15, 19] can characterize surfactant-induced expansion and unfolding of the Trp-cage miniprotein. Therefore, we next analyze the radius of gyration and DSSP to unravel the secondary structure changes.

### 3.3 Structural Unfolding of Trp-cage: Global Expansion and Secondary Structure Loss

The radius of gyration Rg and DSSP secondary-structure maps are capable of distinguishing between increased flexibility and genuine unfolding [15,19]. Both reveal how temperature and surfactant concentration modulate protein compactness and structural stability. At 25 °C, the protein remains relatively compact, consistent with the DSSP map showing a high and persistent α-helical content across most residues, with only limited transitions to turn or coil states (Fig 4 and 5, respectively). Increasing the temperature to 100 °C leads to a noticeable increase in Rg, indicating structural expansion, which correlates with a clear reduction in helical stability and more frequent transitions to turn and coil conformations, particularly at later simulation times (i.e., t>150 ns). A similar trend is observed upon increasing SDS concentration. However, under low SDS, the protein shows early unfolding and a higher Rg at earlier simulation times, suggesting surfactant-assisted destabilization. High SDS conditions appears to promote more loss of the helix, reflected by an increased Rg and a DSSP map dominated by turns and coils, with fragmented and transient helices. This indicates that excessive SDS disrupts native secondary structure and enhances conformational flexibility [3, 7, 8, 15].

**Fig 4.**
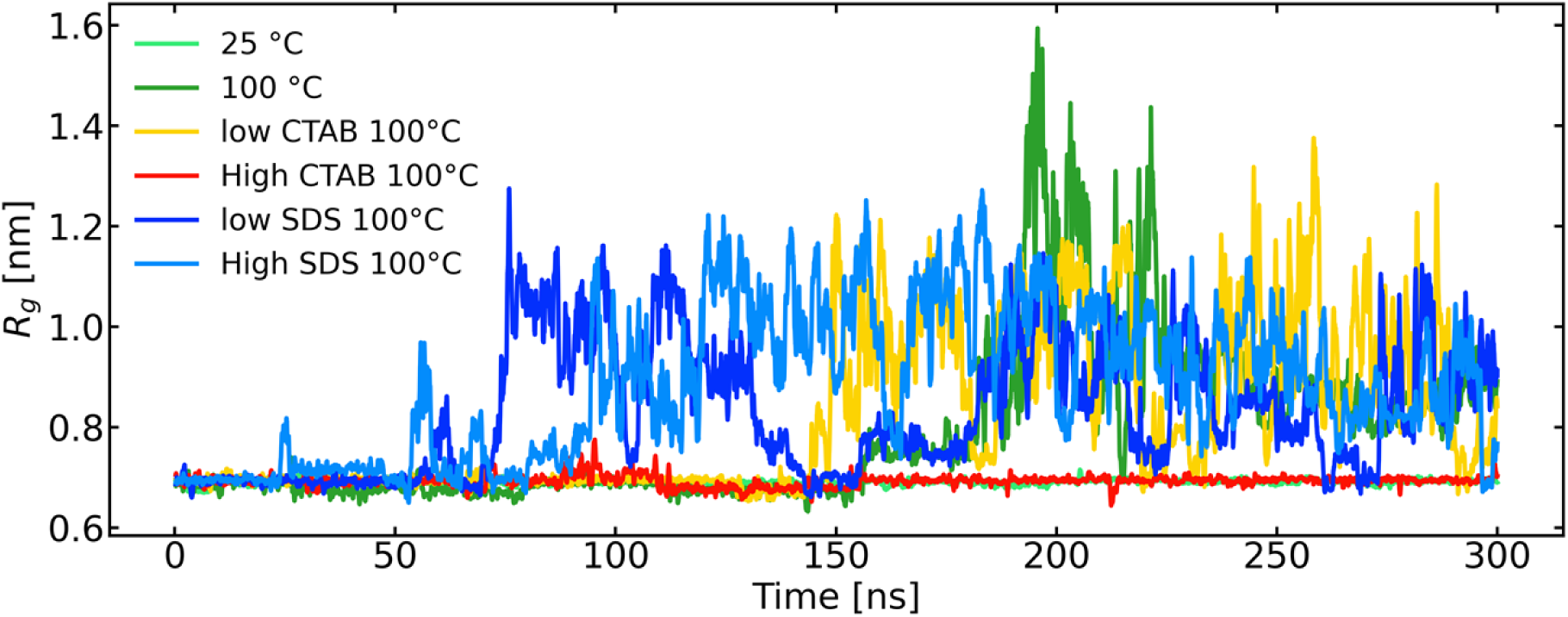
Evolution of the radius of gyration of trp-cage on the simulated timescale (running average).

**Fig 5.**
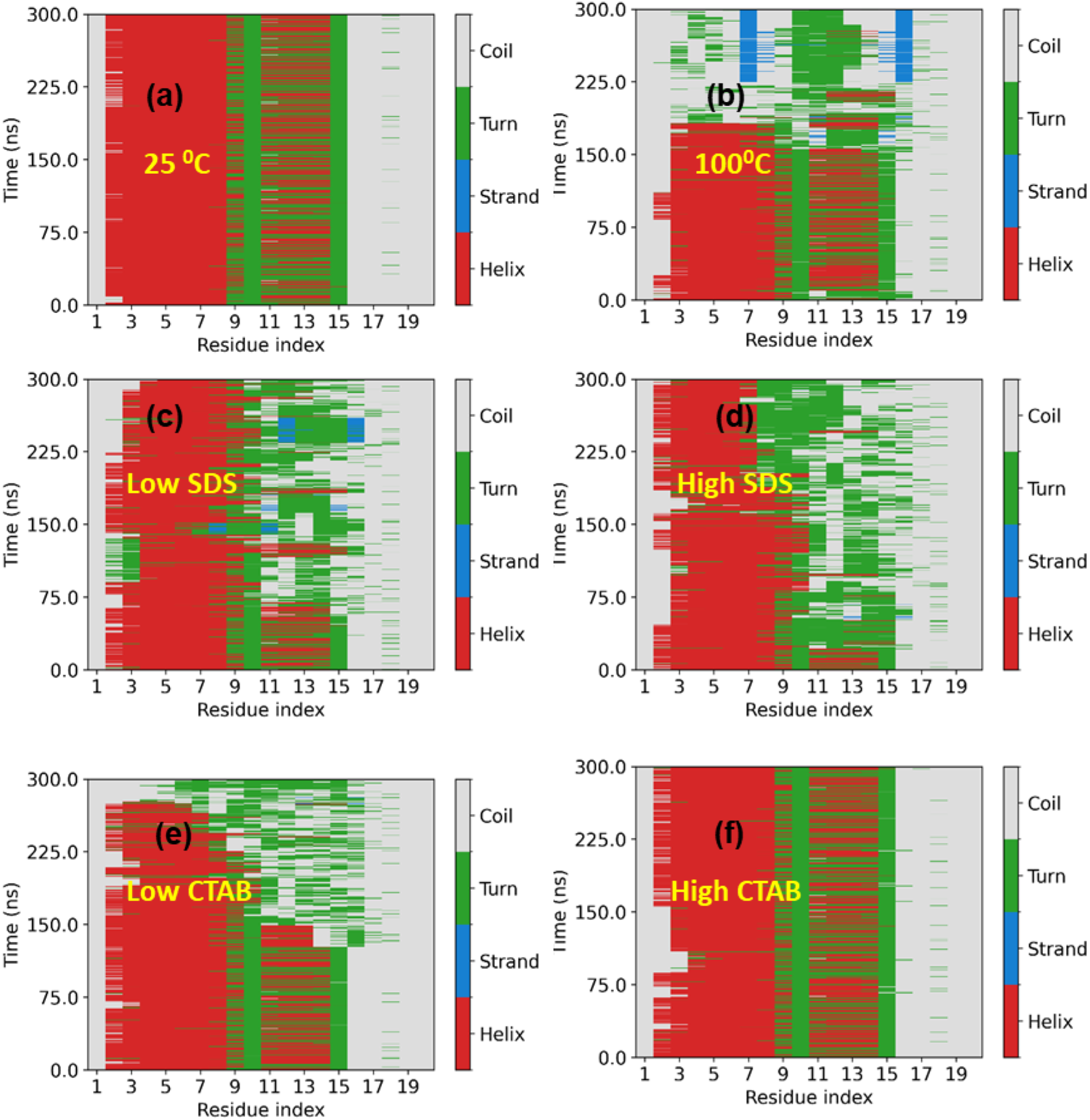
(a-f) DSSP plot of the trp-cage secondary structure.

For CTAB, the protein exhibits stronger helix preservation than in SDS at comparable concentrations. Although some increase in Rg and coil formation is observed at low CTAB, the unfolding rate is slower and delayed than in SDS, suggesting that CTAB stabilizes compact, helical conformations more effectively [8, 19]. Under high CTAB, the Rg remains low and the DSSP plot shows dominant helical content throughout the trajectory. Overall, the combined Rg and DSSP analyses demonstrate that increased temperature and surfactant concentration promote protein expansion and secondary-structure destabilization, with SDS exerting a stronger denaturing effect than CTAB.

While the radius of gyration and secondary-structure analyses provide clear evidence of global expansion and helix destabilization under elevated temperature and surfactant conditions, they do not fully capture how these changes propagate at the level of tertiary packing. In particular, the Trp-cage motif relies on a tightly packed hydrophobic core rather than secondary structure alone for its stability [23]. To determine whether the observed unfolding reflects a true disruption of native tertiary contacts—or merely enhanced backbone flexibility—we next examine the integrity of the hydrophobic core by focusing on tryptophan-centred interactions. Accordingly, The next section probes changes in Trp–hydrophobic residue distances, native contact preservation, and solvent exposure to directly assess collapse or destabilization of the Trp cage under different environmental conditions.

### 3.4 Hydrophobic Core Integrity (Trp-cage collapse)

To probe the integrity of the Trp-cage hydrophobic core, we analyzed both the distribution of the distance between the Trp side chain and core hydrophobic residues, and the time evolution of the hydrophobic core solvent-accessible surface area (SASA).

At 25 °C, the Trp–core distance distribution is sharply peaked around ∼0.25–0.30 nm, indicating a tightly packed hydrophobic core with Trp remaining buried throughout the simulation (Fig 6a) [23]. Increasing the temperature to 100 °C broadens the distribution slightly but retains a dominant peak at short distances, suggesting that thermal fluctuations alone are insufficient to fully disrupt core packing on the simulated timescale.

**Fig 6.**
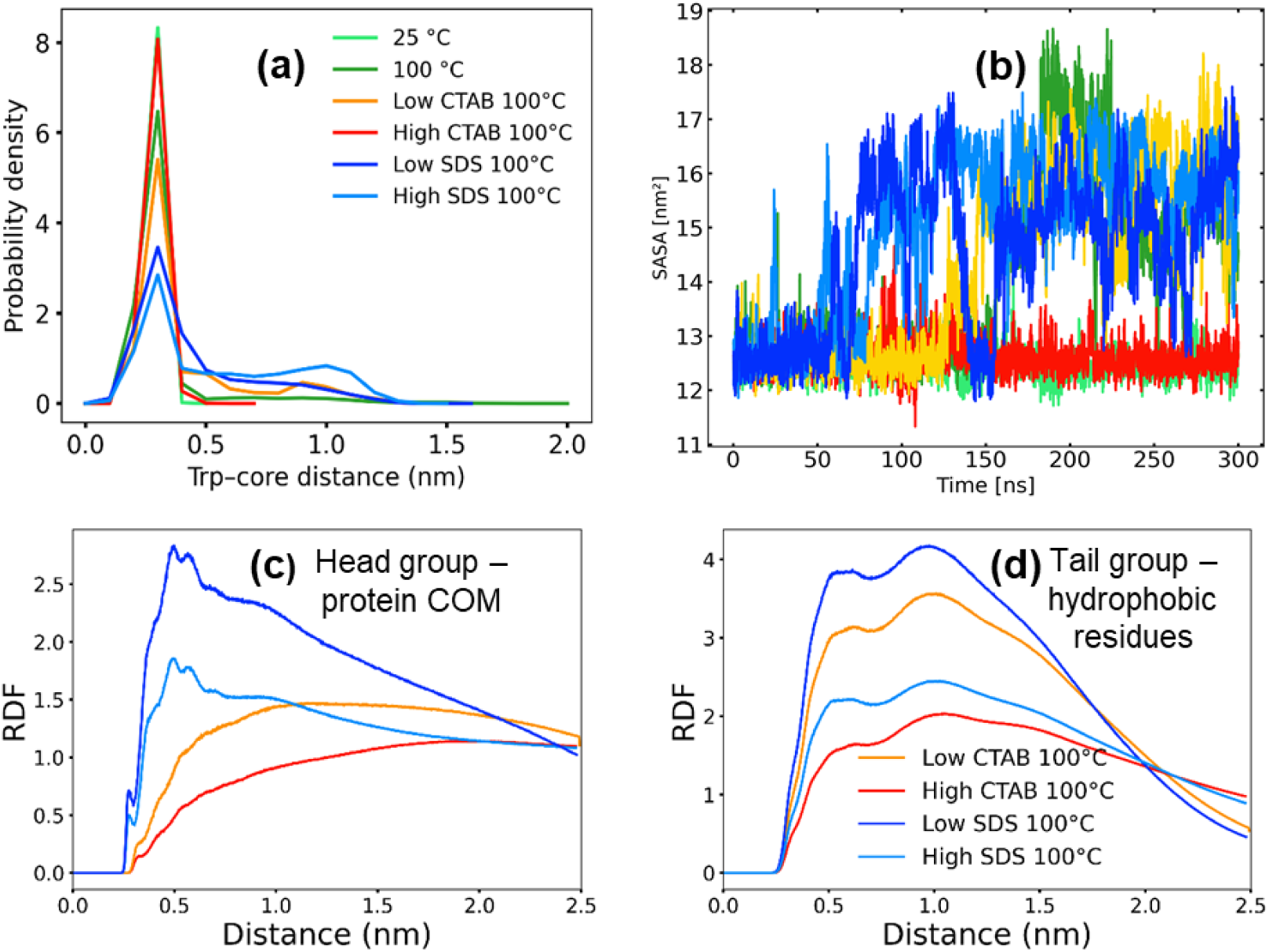
(a) Distribution of the distance between the Trp side chain and core hydrophobic residues (b) hydrophobic core solvent-accessible surface area (c) radial distribution functions of surfactant headgroup and protein center of mas and (d) radial distribution functions of surfactant tailgroup and hydrophobic residues

In contrast, the presence of SDS leads to a lowering of the peak and a shift of the Trp–core distance distribution toward larger values, accompanied by a substantial broadening of the distribution. This effect is concentration dependent, with high SDS exhibiting higher broadening of the distribution, consistent with frequent Trp detachment from the hydrophobic core. These observations indicate a loss of native tertiary contacts and destabilization of the Trp-cage fold in the presence of SDS [3, 15]. In addition, the radial distribution functions reveal a pronounced accumulation of SDS headgroups around the protein canter of mass, indicating strong preferential solvation (Fig 6c). More importantly, SDS alkyl tails exhibit a high and well-defined first coordination peak with hydrophobic protein residues, both at high and low surfactant concentration, consistent with direct tail insertion and micelle-assisted destabilization (Fig 6d).

The SASA analysis supports this interpretation. While the hydrophobic core SASA remains low and stable at 25 °C, SDS systems exhibit a marked increase in core exposure over time, and with less fluctuation at higher surfactant concentration (Fig 6b). In pure water at 100 °C, the increase in SASA is delayed up to 150 ns, consistent with the observed delayed unfolding kinetics. The increased SASA in SDS solutions reflects solvent penetration into the hydrophobic core following Trp displacement, consistent with core opening and partial unfolding.

In comparison, CTAB shows a more modest effect on the Trp–core distances and hydrophobic SASA. Although some broadening of the distance distribution is observed at low CTAB concentration, the dominant short-distance peak is preserved and core SASA remains low and stable up to 125 ns, suggesting delayed unfolding kinetics. Also, the radial distribution functions reveal that CTAB displays weaker headgroup association and broad tail–hydrophobic contact first peak, suggesting a less disruptive interaction mechanism. This suggests that CTAB interactions are largely superficial and do not promote deep penetration into the hydrophobic core, thereby preserving partial tertiary packing even at elevated temperature.

From a practical standpoint, the observed stabilization of Trp-cage by CTAB under high-temperature conditions has important implications for protein formulation science. Thermal denaturation and aggregation represent major obstacles in the storage and global distribution of biologics and research-use-only (RUO) proteins, particularly during transportation from manufacturing facilities to end users, where temperature control may be limited. Our findings suggest that cationic surfactants can mitigate thermally induced unfolding of cationic proteins by preserving hydrophobic core integrity, thereby offering a mechanistic basis for their potential use as stabilizing excipients.

### 3.5 Specific and Cooperative Surfactant Binding

The per-residue contact frequency reveals distinct binding patterns for SDS and CTAB (Fig 7a). SDS exhibits substantially higher contact frequencies across many residues compared to CTAB, particularly at hydrophobic and aromatic positions such as TYR3, ILE4, TRP6, LEU7, and PRO12. These residues are known contributors to the Trp-cage hydrophobic core and surrounding stabilizing contacts [23]. The elevated SDS contacts at these sites indicate deep and persistent association with the protein surface, consistent with SDS’s strong electrostatic anchoring followed by hydrophobic penetration. In contrast, CTAB shows lower and more localized contact frequencies, even at high concentration. Contacts are mainly restricted to exposed residues such as TYR3, PRO12, and ARG16, with relatively weak engagement of core-forming residues (e.g., TRP6 and ILE4). Interestingly, CTAB at high concentration made zero contact with TRP6, further confirming its protective effect against thermal denaturation. This suggests that CTAB primarily interacts superficially, without extensively disrupting the internal packing of the Trp-cage fold. Increasing surfactant concentration leads to similar trends: high SDS strengthens contacts across nearly all residues, whereas high CTAB leads to only modest increases, reinforcing the idea of fundamentally different binding modes rather than a simple concentration effect.

**Fig 7.**
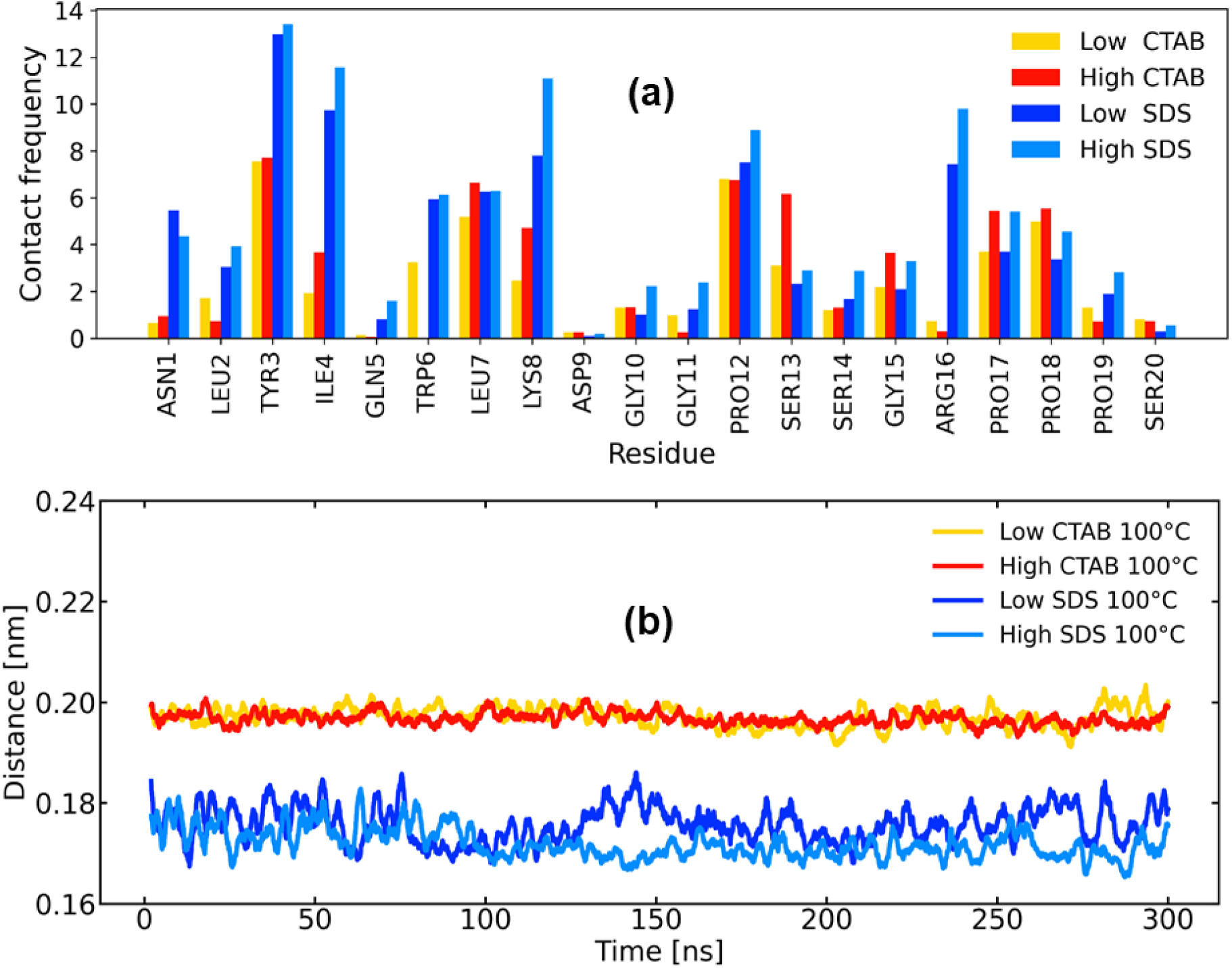
(a) per-residue contact frequency of the surfactants and the protein and (b) global minimum distance between the protein and surfactant molecules.

The global minimum distance between the protein and surfactant molecules (Fig 7b) further supports this interpretation. SDS maintains a consistently shorter protein–surfactant distance (∼0.17–0.18 nm) throughout the simulation, indicating tight and persistent binding. Notably, this close contact is sustained over long timescales, consistent with the formation of stable surfactant–protein complexes that facilitate structural destabilization observed earlier in RMSD, Rg, DSSP, and Trp-core analyses. In contrast, CTAB systems exhibit larger average distances (∼0.19–0.20 nm) with reduced fluctuations, suggesting weaker and more transient interactions. The absence of progressive distance reduction in CTAB simulations implies that CTAB does not progressively invade the protein interior, even at elevated concentration. Together, the residue-specific and global contact analyses demonstrate that SDS binds strongly and pervasively, engaging both surface and core-associated residues, thereby promoting unfolding and hydrophobic core disruption. In contrast, CTAB binds more weakly and selectively, primarily at surface-exposed sites, which limits its ability to destabilize the Trp-cage fold.

These findings provide a direct molecular explanation for the protective effect of high CTAB concentration against thermal denaturation and the enhanced denaturing capability of SDS at low and high concentration observed in earlier structural and tertiary-contact analyses.

### 3.6 Intermediate States and Free Energy Landscape

The conformational behavior of the system under different temperature and surfactant conditions was analyzed by RMSD-based clustering and two-dimensional free-energy surfaces (FES) [15, 19] projected onto the first two collective variables (PC1/RMSD and PC2/Rg) (Fig 8 and 9, respectively). A strong correspondence is observed between the shape of the free-energy landscapes and the resulting cluster population distributions.

**Fig 8.**
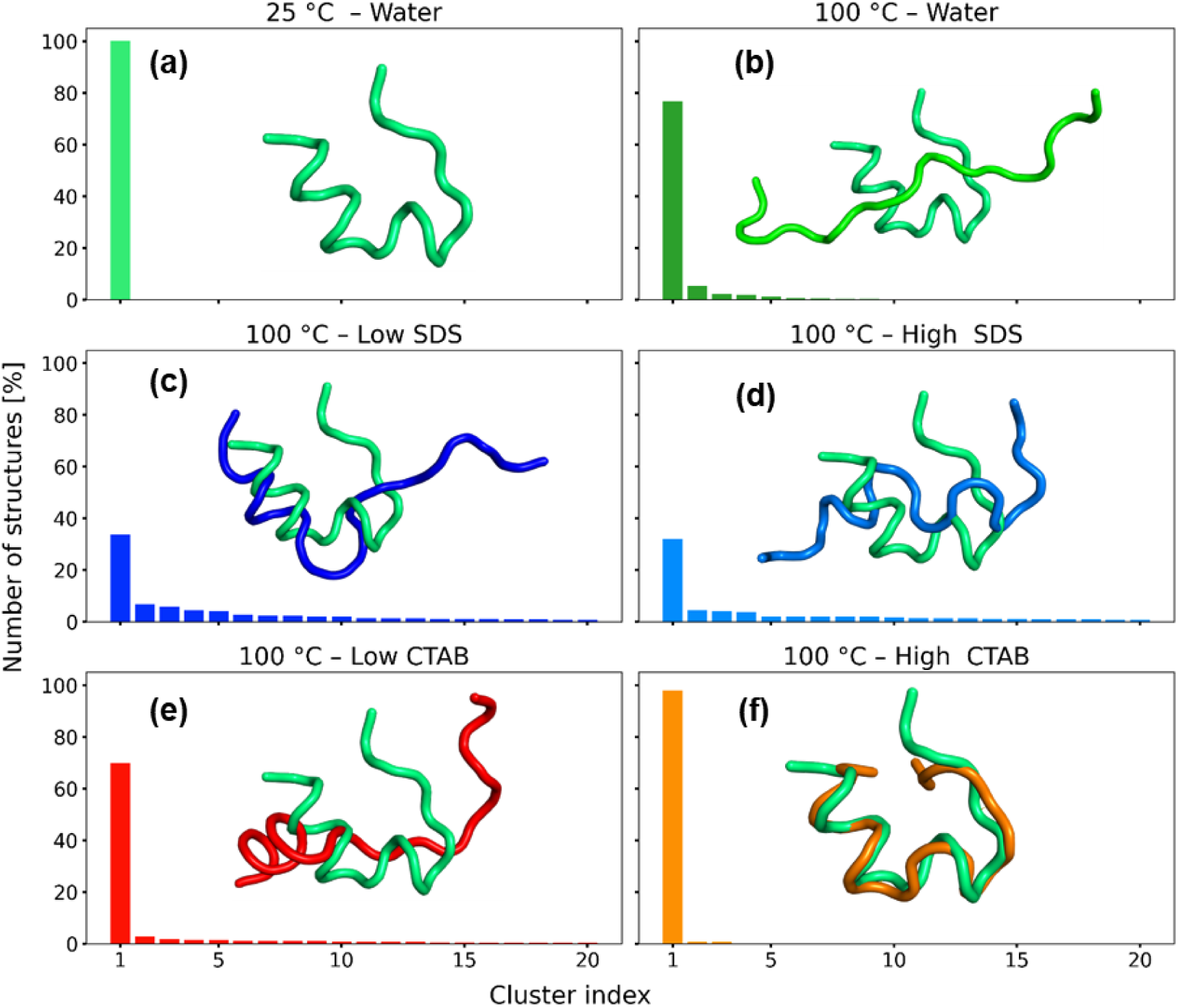
(a-f) clustering analysis of the protein-surfactant interactions. Inset represents the cluster aligned to the reference cluster at 25 °C.

**Fig 9.**
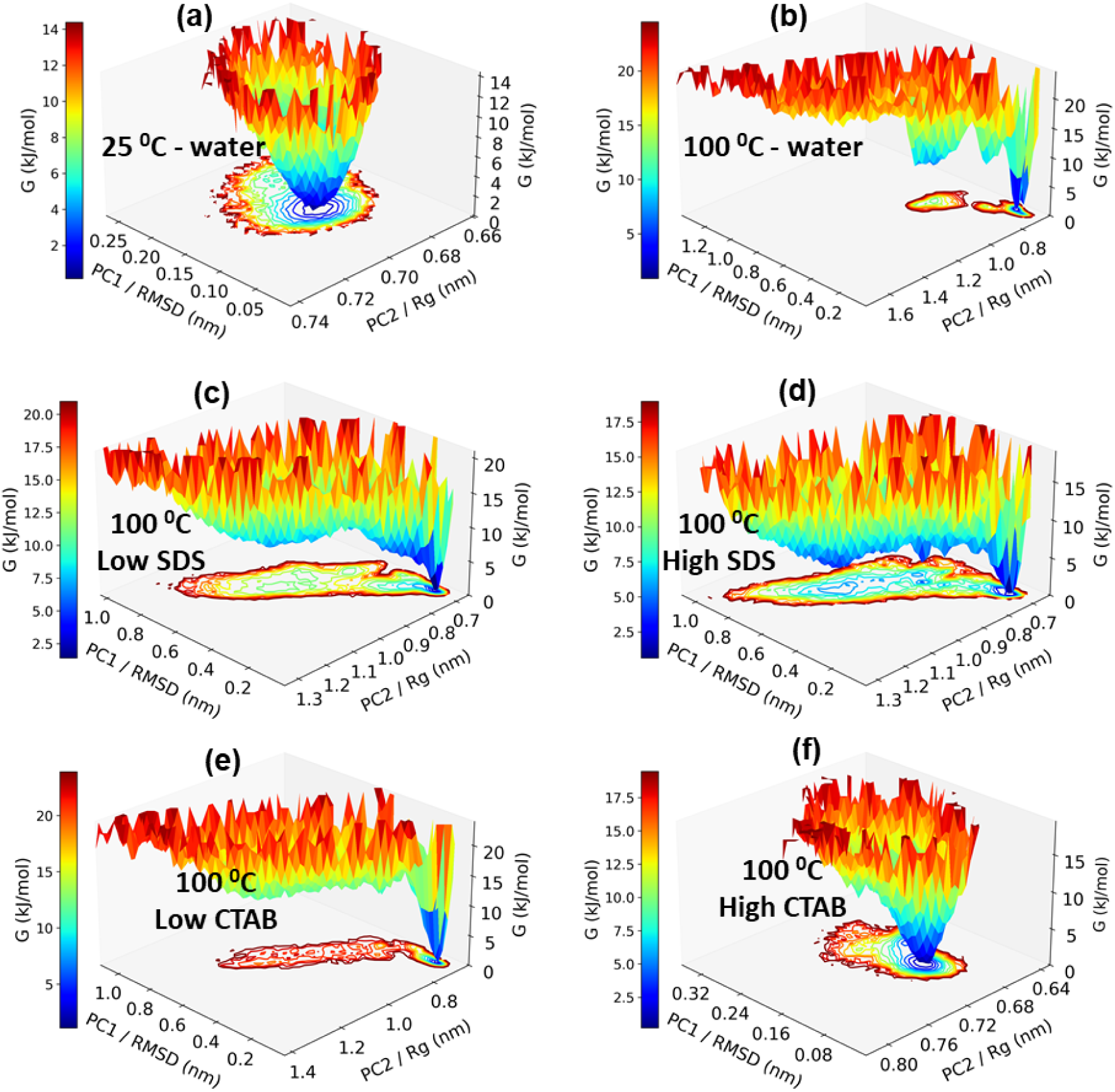
(a-f) free energy landscapes of the protein.

At 25 °C in pure water, the free-energy surface is characterized by a single, deep, and well-defined minimum with a narrow distribution along both collective variables (Fig 9a). This indicates a highly stable conformational ensemble with limited structural fluctuations. Consistent with this observation, clustering analysis identifies a single dominant cluster comprising 100% of the sampled structures, confirming that the system remains confined to one conformational basin under these conditions (Fig 8a). Upon increasing the temperature to 100 °C in pure water, the free-energy landscape becomes significantly distorted and elongated, particularly along the RMSD coordinate. The appearance of multiple shallow minima reflects enhanced conformational flexibility and increased thermal sampling. Correspondingly, the clustering analysis reveals a substantial increase in the number of clusters, with the overlay showing major structural extension with elongated loops and loss of compactness, although a dominant cluster is still present, accounting for approximately 77% of the sampled configurations. This indicates partial retention of a preferred structural state alongside frequent transitions to alternative conformations.

In the presence of SDS at 100 °C, the free-energy landscapes exhibit pronounced ruggedness and an expanded accessible conformational space. Both low and high SDS concentrations lead to fragmentation of the free-energy surface into multiple competing basins. This behavior is mirrored in the clustering results, where the population of the largest cluster decreases markedly (approximately 30–35%), accompanied by a long tail of smaller clusters with non-negligible populations. In addition, the overlay shows a strong deviation from the reference structure. These findings indicate that SDS promotes substantial conformational heterogeneity and destabilization of the dominant folded state at elevated temperature. In contrast, the CTAB systems display concentration-dependent effects. At lower CTAB concentration, the free-energy landscape becomes more elongated and heterogeneous, and the population of the largest cluster decreases to approximately 70%, indicating the emergence of additional metastable states. At higher CTAB concentration and 100 °C, the free-energy surface retains a funnel-like shape with a clearly defined minimum, albeit broader than that observed at 25 °C. Clustering analysis confirms this stabilization, with nearly all structures (≈98%) belonging to a single dominant cluster and the overlay showing close overlaps.

Overall, these results demonstrate that temperature is the primary driver of conformational diversification, while surfactants modulate this effect in a concentration-dependent manner. Systems exhibiting deep and localized free-energy minima correspond to highly populated single clusters, whereas rugged and extended landscapes give rise to broad cluster population distributions. SDS enhances conformational heterogeneity, whereas CTAB partially counteracts thermal destabilization, particularly at high concentration.

## 4. Conclusion

In this study, we employed all-atom molecular dynamics simulations to elucidate the effects of temperature and surfactant chemistry on protein conformational stability. Free-energy landscapes derived from principal component analysis reveal that increasing temperature leads to a pronounced expansion of the accessible conformational space, indicating partial unfolding and enhanced structural heterogeneity. This effect is further modulated by the presence of surfactants in a charge and concentration-dependent manner.

Cluster population analysis demonstrates that SDS strongly destabilizes the protein at elevated temperature, as evidenced by the redistribution of structural populations across multiple low-occupancy clusters and the loss of a dominant native-like state. In contrast, CTAB preserves a higher degree of conformational order, particularly at higher concentrations, where a small number of clusters continue to dominate the ensemble. These findings indicate fundamentally different mechanisms of protein–surfactant interaction for anionic and cationic surfactants.

Radial distribution function analysis provides molecular insight into these differences. SDS exhibits strong headgroup accumulation near the protein surface and significant penetration of its hydrophobic tails into protein hydrophobic regions, consistent with micelle-assisted unfolding driven by hydrophobic interactions. CTAB, by contrast, shows weaker headgroup association and reduced tail–hydrophobic residue contacts, correlating with its comparatively stabilizing effect.

Overall, our results highlight the central role of surfactant tail–protein hydrophobic interactions in governing protein destabilization at high temperature, while emphasizing that electrostatic effects alone are insufficient to explain the observed conformational changes. This work advances a mechanistic understanding of surfactant-induced protein unfolding and provides a framework for rationally tuning solvent and surfactant environments to control protein stability under extreme conditions.

## Contributions

O.S. N. and H.B conceived and designed the simulation and O.S.N performed the simulation and wrote the manuscript. O.S. N., H.B and U.E.O reviewed the manuscript.

